# S-Nitrosylation of STIM1 by neuronal nitric oxide synthase inhibits store-operated Ca^2+^ entry

**DOI:** 10.1101/304022

**Authors:** Le Gui, Jinhui Zhu, Xiangru Lu, Stephen M. Sims, Wei-Yang Lu, Peter B. Stathopulos, Qingping Feng

## Abstract

Store-operated Ca^2+^ entry (SOCE) mediated by stromal interacting molecule-1 (STIM1) and Orai1 represents a major route of Ca^2+^ entry in mammalian cells and is initiated by STIM1 oligomerization in the endoplasmic or sarcoplasmic reticulum (ER/SR). However, the effects of nitric oxide (NO) on STIM1 function are unknown. Neuronal NO synthase (nNOS) is located in the SR of cardiomyocytes. Here, we show that STIM1 is susceptible to S-nitrosylation. nNOS deficiency or inhibition enhanced Ca^2+^ release-activated Ca^2+^ channel current (*I_CRAC_*) and SOCE in cardiomyocytes. Consistently, NO donor S-nitrosoglutathione (GSNO) inhibited STIM1 puncta formation and *I_CRAC_* in HEK293 cells, but this effect was absent in cells expressing the Cys49Ser/Cys56Ser STIM1 double mutant. Furthermore, NO donors caused Cys49 and Cys56-specific structural changes associated with reduced protein backbone mobility, increased thermal stability and suppressed Ca^2+^-depletion-dependent oligomerization of the luminal Ca^2+^-sensing region of STIM1. Collectively, our data show that S-nitrosylation of STIM1 suppresses oligomerization *via* enhanced luminal domain stability and rigidity, and inhibits SOCE in cardiomyocytes.

## INTRODUCTION

Store-operated Ca^2+^ entry (SOCE) is the main mechanism of Ca^2+^ influx into the non-excitable cells and regulates Ca^2+^ homeostasis in excitable cells including neurons, skeletal muscle cells and cardiomyocytes [1–3]. A major function of SOCE is to refill the depleted sarcoplasmic/endoplasmic reticulum (SR/ER) Ca^2+^ stores via elevations in cytosolic Ca^2+^ sourced from the extracellular space. Increases in cytosolic Ca^2+^ by SOCE also regulate various cellular functions including exocytosis, muscle contraction, cell mobility, gene transcription, cell proliferation and apoptosis [3]. Stromal interaction molecules (STIM1 and 2) and Orai family members (Orai1-3) are proteins involved in the activation and formation of Ca^2+^ release activated Ca^2+^ (CRAC) channels, which mediate a highly Ca^2+^-selective conductance (*I_CRAC_*) and represent a major SOCE pathway [3]. STIM1 is a single pass transmembrane protein that is primarily localized to the SR/ER membrane. The EF-hand and the sterile alpha motif (SAM) domains function to sense decreases in SR/ER luminal Ca^2+^ levels through a mechanism which couples a destabilization of the EF-hand:SAM domain interface with dimerization and oligomerization [4–6]. Moreover, the oligomerization of this EF-SAM region induces a conformational rearrangement on the cytosolic portion of STIM1 that culminates in translocation of STIM1 molecules to SR/ER-plasma membrane junctions where Orai1 subunits are recruited, assembled into channels and gated to permit Ca^2+^ entry into the cytosol [7–9]. Apart from SR/ER Ca^2+^ levels, STIM1 is sensitive to transient temperature increases such that cooling allows heat-activated STIM1 to couple with Orai-formed channels [10]. Additionally, post-translational modifications including phosphorylation, *N*-glycosylation and *O*-GlcNAcylation, regulate STIM1-Orai1 coupling and SOCE activation [10–13]. However, it is not known if STIM1 function is regulated by S-nitrosylation.

S-Nitrosylation is a reversible, covalent addition of the NO moiety to cysteine thiols forming an S-nitrosoprotein (SNO-protein) and is one of the principal protein post-translational modifications that regulate protein function and cell signalling [14]. S-Nitrosylation can occur when a thiol group reacts with NO in the presence of an electron acceptor. This reaction can be carried out by protein nitrosylases, which include metal-to-Cys and Cys-to-Cys nitrosylases. Over the past decade, more than 1,000 proteins, which regulate diverse cellular and organ functions, have been identified to be susceptible to S-nitrosylation [14]. Most of the greater than 50,000 proteins in the human proteome have not been assessed for S-nitrosylation, and importantly, the physiological consequences of this post-translational modification remain to be determined. Endothelial and neuronal NOS (eNOS and nNOS) are expressed in the sarcolemma and SR of cardiomyocytes, respectively. It has been shown that eNOS induces S-nitrosylation of G protein-coupled receptor kinase 2 (GRK2), β-arrestin 2 and dynamin, while nNOS S-nitrosylates Ca^2+^ handling proteins including the L-type Ca^2+^ channel, RyR2 and SERCA2a in cardiomyocytes [14–16].

The luminal segment of STIM1 preceding the EF-SAM domain contains two cysteine residues (*i.e.* Cys49 and Cys56) which are conserved from *Drosophila* to humans [17]. In response to oxidative stress, Cys56 undergoes S-glutathionylation, activating STIM1 and CRAC channels in DT40 lymphocytes [18]. To date, it is not known if these cysteine residues are susceptible to S-nitrosylation by NO or whether S-nitrosylation of STIM1 regulates CRAC channel activity in mammalian cells. Here, using primary nNOS^−/−^ cardiomyocyte cell culture, HEK293 cell lines and recombinantly isolated luminal STIM1 domains we show that nNOS inhibits *I_CRAC_* and SOCE via S-nitrosylation of Cys49 and Cys56, which increases the rigidity of the protein backbone, stabilizes the Ca^2+^ sensing domain and inhibits oligomerization. This suppressive effect is opposite to S-glutathionylation [18] and highlights how post-translational modifications can fine-tune the Ca^2+^ sensing response of STIM1 and SOCE commensurate with a changing cellular environment.

## RESULTS

### STIM1 S-nitrosylation in the heart

To assess STIM1 S-nitrosylation, neonatal mouse hearts were isolated and homogenized. S-Nitrosylated proteins were biotin-tagged with the biotin-switch protocol using ascorbic acid to selectively reduce S-NO groups. S-Nitrosylated STIM1 was subsequently detected by western blotting. Our results show that relative levels of detectable S-nitrosylated STIM1 proteins were lowered by 49% in nNOS^−/−^ compared to WT hearts (*P*<0.01, **Fig. 1a**). Since disulfide bonds can form as a consequence of S-nitrosylation [19], we assessed the propensity for intramolecular disulfide formation using a gel migration assay previously used for STIM1 [20]. STIM1 from WT hearts exhibited an increased mobility under non-reducing compared to reducing conditions, while STIM1 from nNOS^−/−^ or eNOS^−/−^ hearts showed no change in mobility (**Fig. S1**). The data is consistent with the relatively higher levels of S-nitrosylation observed for WT compared to nNOS^−/−^ samples (**Fig. 1a**). Interestingly, deficiency in eNOS lowered STIM1 S-nitrosylation by 52% (**Fig. S2**), suggesting eNOS also causes STIM1 S-nitrosylation in the heart. S-Nitrosylation of isolated human STIM1 luminal domain (residues 23-213) protein by NO donor S-nitrosoglutathione (GSNO) was demonstrated using a fluorescein-5-maleimide labeling assay (**Fig. S3**).

**Figure 1.**
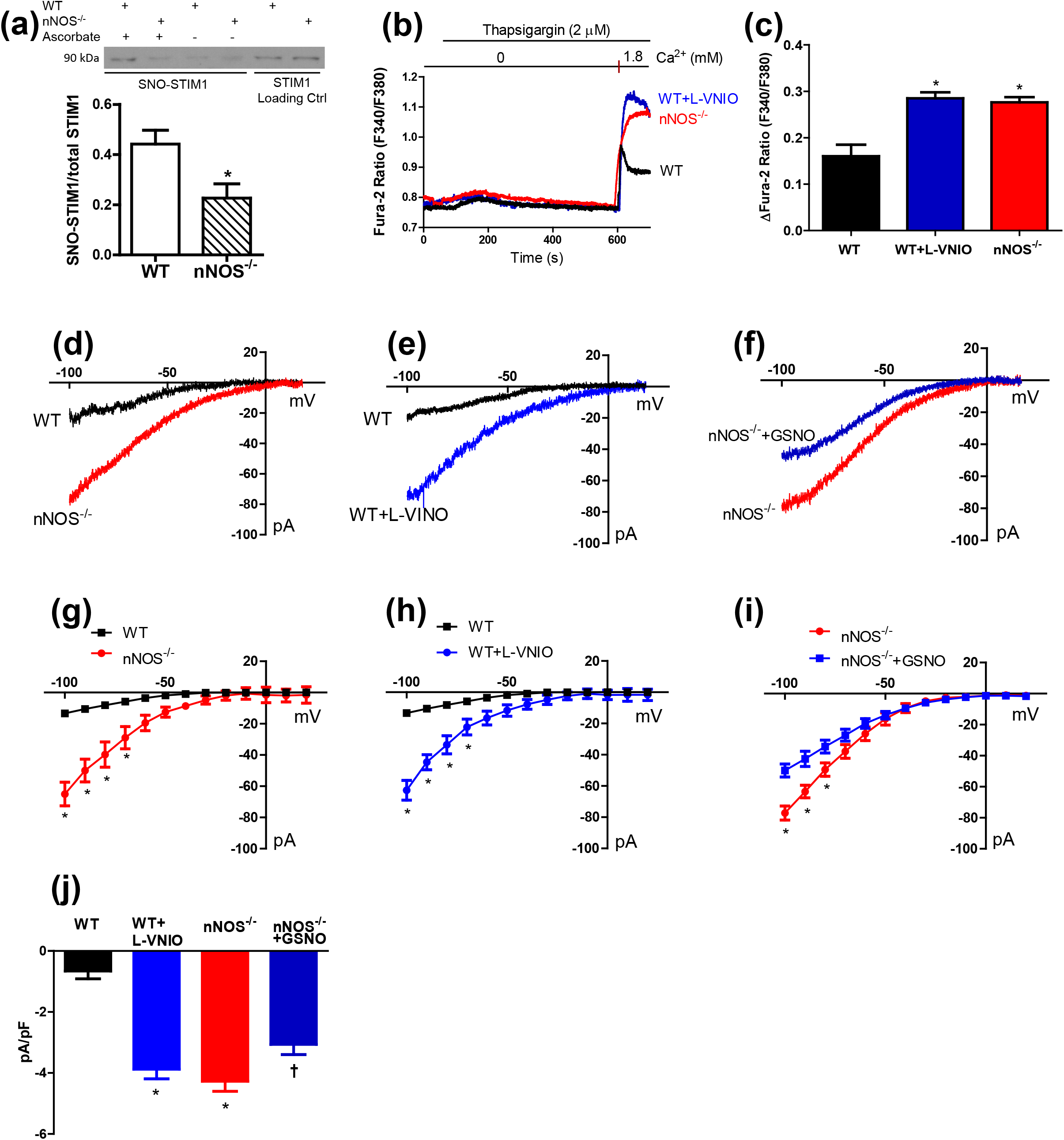
Deficiency in nNOS decreases S-nitrosylation of STIM1 and increases SOCE *I_CRAC_* in cardiomyocytes. **(a)** S-Nitrosylation of STIM1 was assessed by iodoTMT labeling followed by Western blotting in WT and nNOS^−/−^ neonatal hearts with total STIM1 served as a loading control. Upper panel is a representative blot. \**P*<0.05 vs. WT, n=4 hearts per group. **(b)** Representative traces of Fura-2 ratios indicating activation of SOCE in neonatal cardiomyocytes. **(c)** Quantification of Fura-2 ratio changes in WT, WT+L-VNIO (100 μM), and nNOS^−/−^ cardiomyocytes in response to restoration of extracellular Ca^2+^ (1.8 mM). N = 18-19 cells per group from 3-4 independent cell cultures. \**P*□ 0.01 vs. WT. **(d-f)** Representative traces of *I_CRAC_* in neonatal cardiomyocytes. **(g-i)** Summary of current-voltage relationship of *I_CRAC_* in WT and nNOS^−/−^ cardiomyocytes with or without L-VNIO (100 μM) or NO donor S-nitrosoglutathione (GSNO, 250 μM). (**j**) Current density to whole-cell capacitance at −80 mV. N = 8-9 cells per group from 4 independent cell cultures. \**P*□ 0.01 vs. WT or corresponding controls for c and g-i. †*P*<0.05 vs. nNOS^−/−^ cardiomyocytes.

### SOCE and *I_CRAC_* are increased in nNOS^−/−^ neonatal cardiomyocytes

Since nNOS is the predominant NOS isoform in cardiomyocytes and is localized to the SR in close proximity with STIM1, the present study is focused on the role of nNOS in regulating SOCE and *I_CRAC_*. To this end, cultured neonatal cardiomyocytes were employed to analyze Ca^2+^ influx using fura-2. Baseline assessments show similar intracellular Ca^2+^ levels among WT, WT+L-VNIO and nNOS^−/−^ cardiomyocytes with fura-2 ratios of 0.62±0.02, 0.68±0.03 and 0.67±0.04, respectively (*P*=n.s.). Cells were then incubated in Ca^2+^-free media and treated with thapsigargin (2 μM), an SR Ca^2+^ ATPase (SERCA) inhibitor, for 10 min to deplete SR Ca^2+^ stores. This treatment was followed by incubation with 1.8 mM extracellular Ca^2+^, and the resulting Ca^2+^ influx in cardiomyocytes was recorded using fura-2 fluorescence (**Fig. 1b**). SOCE was significantly higher in nNOS^−/−^ compared to wild-type (WT) cardiomyocytes (*P*<0.05, **Fig. 1c**). Treatment with the nNOS inhibitor N^5^-(1-Imino-3-butenyl)-L-ornithine (L-VNIO, 100 μM) also increased SOCE in WT cardiomyocytes (P<0.05, **Fig. 1b and c**).

The CRAC channels in cultured neonatal cardiomyocytes were primed by excluding Ca^2+^ from the extracellular solution which contained the voltage-gated Ca^2+^ channel blocker verapamil (10 μM). Using whole-cell voltage-clamp recording (intracellular solution contained 10 mM BAPTA, a Ca^2+^ chelator), transmembrane conductance and the current-voltage (I-V) relationship of cultured cardiomyocytes were revealed by a voltage-ramp from −100 to +20 mV in the absence and presence of a CRAC channel blocker BTP2 (20 μM). The BTP2-sensitive component, which reflects *I_CRAC_*, recorded from individual cells was obtained. After adding 1.8 mM Ca^2+^ to the extracellular solution, the I-V curve of cardiomyocytes exhibited as an inwardly rectified conductance, a characteristic feature of *I_CRAC_* (**Fig. 1d-f**). Interestingly, the amplitude of *I_CRAC_* was significantly larger in nNOS^−/−^ cells and L-VNIO treated WT cells in comparison to WT control cells (*P*<0.01, **Fig. 1g and h**). Notably, the NO-donor GSNO (250 μM) significantly inhibited *I_CRAC_* in nNOS^−/−^ cells (*P*<0.05, **Fig. 1f and i**). To control for variability in cell size, current density (pA/pF) at −80 mV is shown in **Fig. 1j**. Together, these results indicate that NO inhibits *I_CRAC_*

### NO-mediated *I_CRAC_* inhibition is dependent on Cys49 and Cys56 of STIM1

Since luminal STIM1 Cys56 is susceptible to modification by oxidative stress [18,20], we hypothesized that the luminal Cys residues are also targets of NO-mediated modification and inhibition of *I_CRAC_*. Thus, we generated single or double mutations (i.e. Cys49Ser, Cys56Ser or Cys49Ser/Cys56Ser) into a full length monomeric cherry fluorescent protein-tagged human STIM1 (mCh-STIM1) construct. WT, single or double cysteine mutants of mCh-STIM1 were transfected into HEK293 cells stably expressing yellow fluorescence protein (YFP)-Orai1[21]. Expression of mCh-STIM1 and YFP-Orai1 in HEK293 cells was confirmed by red and yellow fluorescence signals, respectively. HEK293 cells express little or no nNOS or eNOS as these proteins were not detectable by western blot analysis (**Fig. S4**). Whole-cell voltage-clamp recordings revealed that HEK293 cells stably expressing YFP-Orai1 alone had no detectable BTP2-senstive *I_CRAC_* (**Fig. S5**). In contrast, transfection of WT full length mCh-STIM1 into the YFP-Orai1 expressing HEK293 cells resulted in a BTP2-sensitive *I_CRAC_*, which was inhibited by ∼45% in the presence of GSNO (**Fig. 2a**). Remarkably, the inhibitory effect of GSNO was attenuated to 16% and 9% in cells expressing Cys49Ser or Cys56Ser mCh-STIM1, respectively (**Fig. 2b and c**). Importantly, the effect of GSNO on *I_CRAC_* was completely abolished in cells expressing the Cys49Ser/Cys56Ser double mutant mCh-STIM1 (**Fig. 2d**). The *I_CRAC_* response in the present of GSNO was significantly different between WT and double mutant transfected cells while the baseline currents were similar between the two groups (**Fig. 2e**). These data suggest that Cys49 and Cys56 are likely targets of S-nitrosylation by NO.

**Figure 2.**
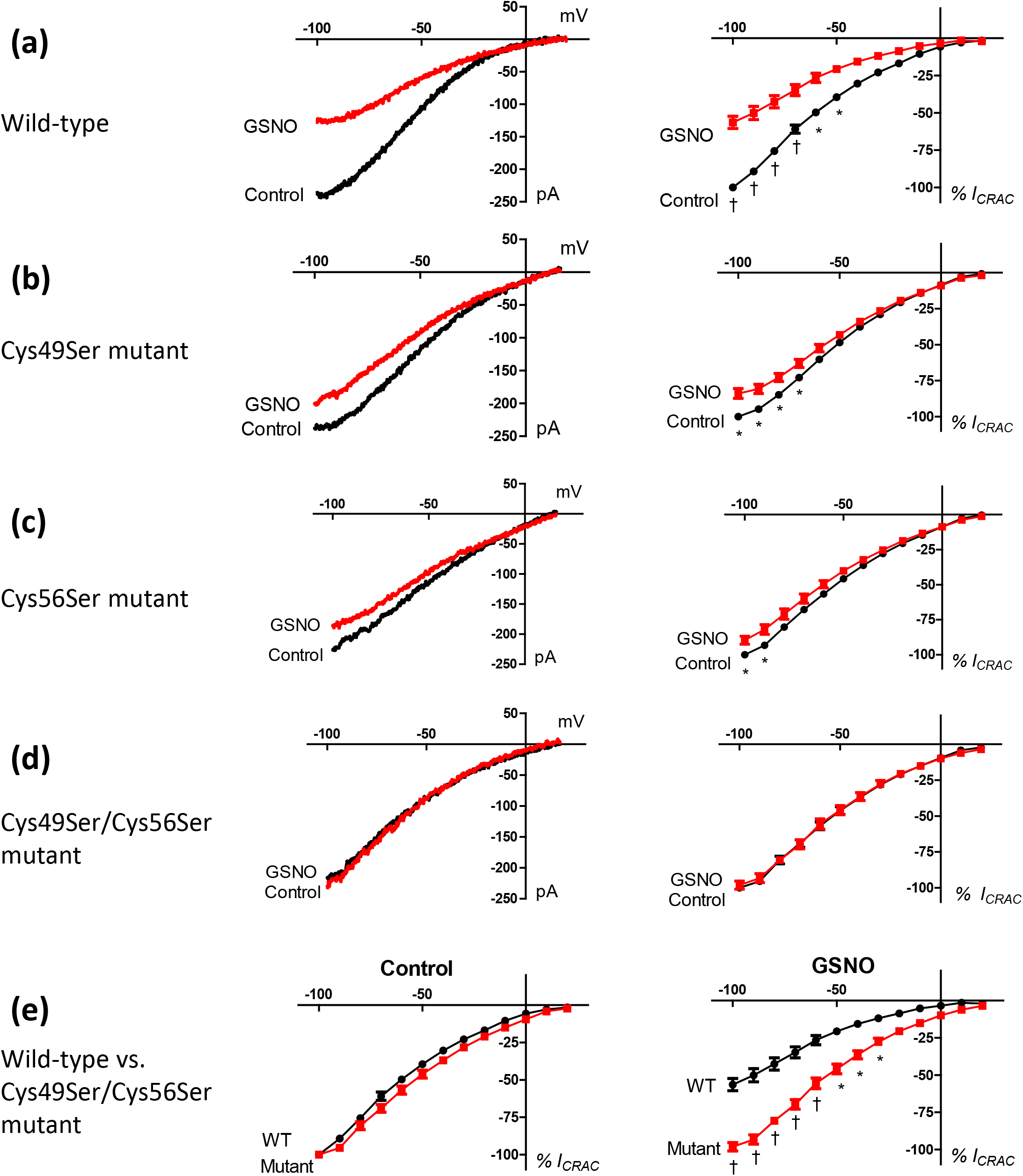
Effects of S-nitrosoglutathione (GSNO) on current-voltage relationship of *I_CRAC_* in full-length wild-type (WT) and STIM1 mutant transfected HEK293 cells stably expressing Orai1. GSNO (250 μM) inhibited *I_CRAC_* of WT STIM1 transfected cells by 45%. **(a)**. This effect was reduced to 16% and 9% in Cys49Ser **(b)** and Cys56Ser STIM1 **(c)** mutant transfected cells, respectively, and was completely abolished in Cys49Ser/Cys56Ser STIM1 double mutant transfected cells **(d)**. Left panels are representative traces. Right panels show summaries of *I_CRAC_*. **(e)** Comparison between WT and Cys49Ser/Cys56Ser STIM1 double mutant transfected cells with and without GSNO (250 μM) treatment. Data are mean ±SEM. N=7-12 cells from 3-4 experiments per group, \**P*□ 0.05, †*P*<0.01 vs control or WT.

### NO rigidifies the non-conserved N-terminal region of STIM1

To confirm NO donors were acting on the Cys49 and Cys56 residues of STIM1, we recombinantly expressed and purified the short N-terminal STIM1 fragment corresponding to residues 24-57 harboring the Cys targets. An ^1^H-^15^N) solution NMR spectrum showed the amide H(N) crosspeaks clustered in the ∼7.5-8.5 ppm region in the presence of 2 mM GSNO and numerous chemical shift differences compared to the spectrum acquired with 2 mM DTT (**Fig. 3a**), indicative of residue-specific structural changes associated with the GSNO-induced modification. We sequentially assigned ∼85 and 84 % of the Cα, Cβ, H, N backbone chemical shifts for the STIM1 24-57 fragment in 2 mM GSNO or 2 mM DTT, respectively, using HNCACB experiments. The H(N) chemical shift perturbation (CSP) map revealed that the largest chemical shift changes were clustered near the Cys49 and Cys56 residues, as expected for Cys-specific S-nitrosylation (**Fig. 3b**). The largest carbon shift changes were localized to the Cα and Cβ atoms of Cys49 and Cys56, consistent with modification of the proximal sulfhydryl groups (**Fig. 3c**). The chemical shift index (CSI) [22] of STIM1 24-57 calculated from the backbone chemical shifts showed that, in isolation, this region does not form regular secondary structure either with or without the NO donor. However, the random coil index (RCI) [23] revealed strikingly higher order parameters (S^2^) for residues clustered near the Cys49 site (**Fig. 3d**), suggesting decreased mobility in the presence of GSNO compared to DTT. Thus, GSNO modifies the Cys49 and Cys56 residues, causing structural changes associated with rigidification of the residues localized around Cys49.

**Figure 3.**
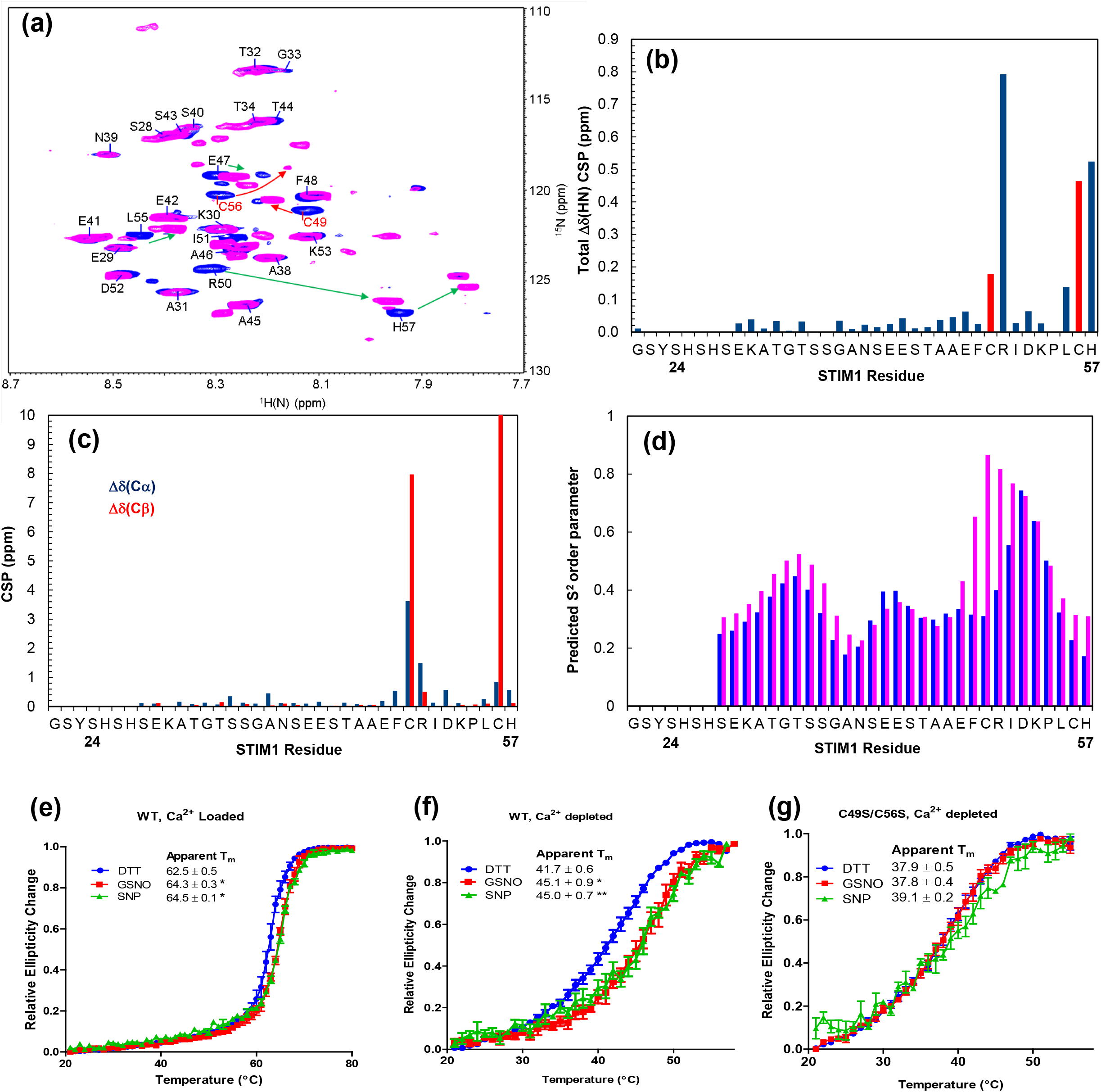
Effects of S-nitrosoglutathione (GSNO) on STIM1 24-57 protein structure and STIM1 23-213 thermal stability. **(a)** ^1^H-^15^N-HSQC spectral overlay of STIM1 24-57 in 2 mM GSNO (magenta) or 2 mM DTT (blue). **(b)** NMR chemical shift perturbation (CSP) of STIM1 57 protein caused by GSNO for the alpha carbon 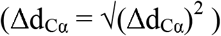 and the beta carbon 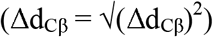. **(c)** Total H(N) 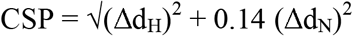 (Δd_N_)^2^ of STIM1 24-47 caused by GSNO. **(d)** Model-free order parameter (S^2^) predicated from the backbone chemical shifts of the GSNO or DTT treated samples. **(e-g)** Thermal stability was assessed by changes in far-UV CD signal of unmodified (1 mM DTT) and S-nitrosylated (1 mM GSNO or 2 mM SNP) STIM1 23213 protein. **(e)** Wild-type, Ca^2+^-loaded. **(f)** Wild-type, Ca^2+^-depleted. **(g)** Cys49Ser/Cys56Ser double mutant, Ca^2+^-depleted. Data are means ± SEM from 3 – 6 separate protein samples. \**P*□ 0.05, \*\**P*<0.01 vs DTT control.

### NO stabilizes the STIM1 luminal domain in a Cys49 and Cys56-dependent manner

We next sought to determine how the NO donor-induced structural and dynamical changes in STIM1 24-57 affect the stability of the entire luminal domain of STIM1. Thus, we assessed stability of the STIM1 23-213 protein by monitoring changes in far-UV circular dichroism (CD) signal as a function of temperature in the presence and absence of NO donors. In the absence of NO donors, the Ca^2+^-loaded STIM1 23-213 protein unfolded with a midpoint temperature of denaturation (T_m_) of 62.5 ± 0.5 °C. Remarkably, either excess GSNO or sodium nitroprusside (SNP) NO donors significantly increased the thermal stability of the Ca^2+^ loaded STIM1 23-213 region by ∼2 °C (**Fig. 3e**). In the absence of NO, the Ca^2+^-depleted luminal domain of STIM1 was highly destabilized compared to the Ca^2+^-loaded state with a T_m_ of 41.7 ± 0.6 °C, as previously demonstrated [24]. While the stabilization observed for the Ca^2+^-loaded protein was a modest ∼2 °C in the presence of the NO donors, the Ca^2+^-depleted STIM1 23-213 protein showed a more marked ∼4 °C increase in T_m_ in the presence of excess GSNO or SNP (**Fig. 3f**). The Cys49Ser/Cys56Ser double mutation of STIM1 23-213 caused an overall destabilization of the protein; however, the T_m_ values for both the Ca^2+^-loaded and Ca^2+^-depleted states were not significantly altered by the NO donors (**Fig. 3g**). Thus, NO significantly enhances the stability of the luminal domain of STIM1 in both the presence and absence of Ca^2+^ in a Cys49- and Cys56-dependent manner.

### NO inhibits oligomerization of the STIM1 luminal domain in Cys49 and Cys56-dependent manner

Ca^2+^ depletion induced destabilization-coupled oligomerization initiates STIM1 activation and SOCE [6–8,25–27]. Using dynamic light scattering (DLS), Ca^2+^-depleted WT STIM1 23-213 showed an earlier decay in the normalized autocorrelation function when treated with GSNO compared to DTT, indicative of smaller particle sizes. GSNO did not elicit a similar shift for the Cys49Ser/Cys56Ser protein (**Fig. 4a**). The average monodisperse particle size weighted toward the largest particle was significantly smaller for the WT protein in the presence of GSNO compared to DTT, while the NO donor did not similarly affect the Cys49Ser/Cys56Ser protein (**Fig. 4b**). Regularized deconvolution of the polydisperse sizes showed a distinct distribution of ∼monomer/dimer hydrodynamic radii for WT protein in the presence of the NO donor that was absent with DTT treatment and with the Cys49Ser/Cys56Ser protein (**Fig. 4c**).

**Figure 4.**
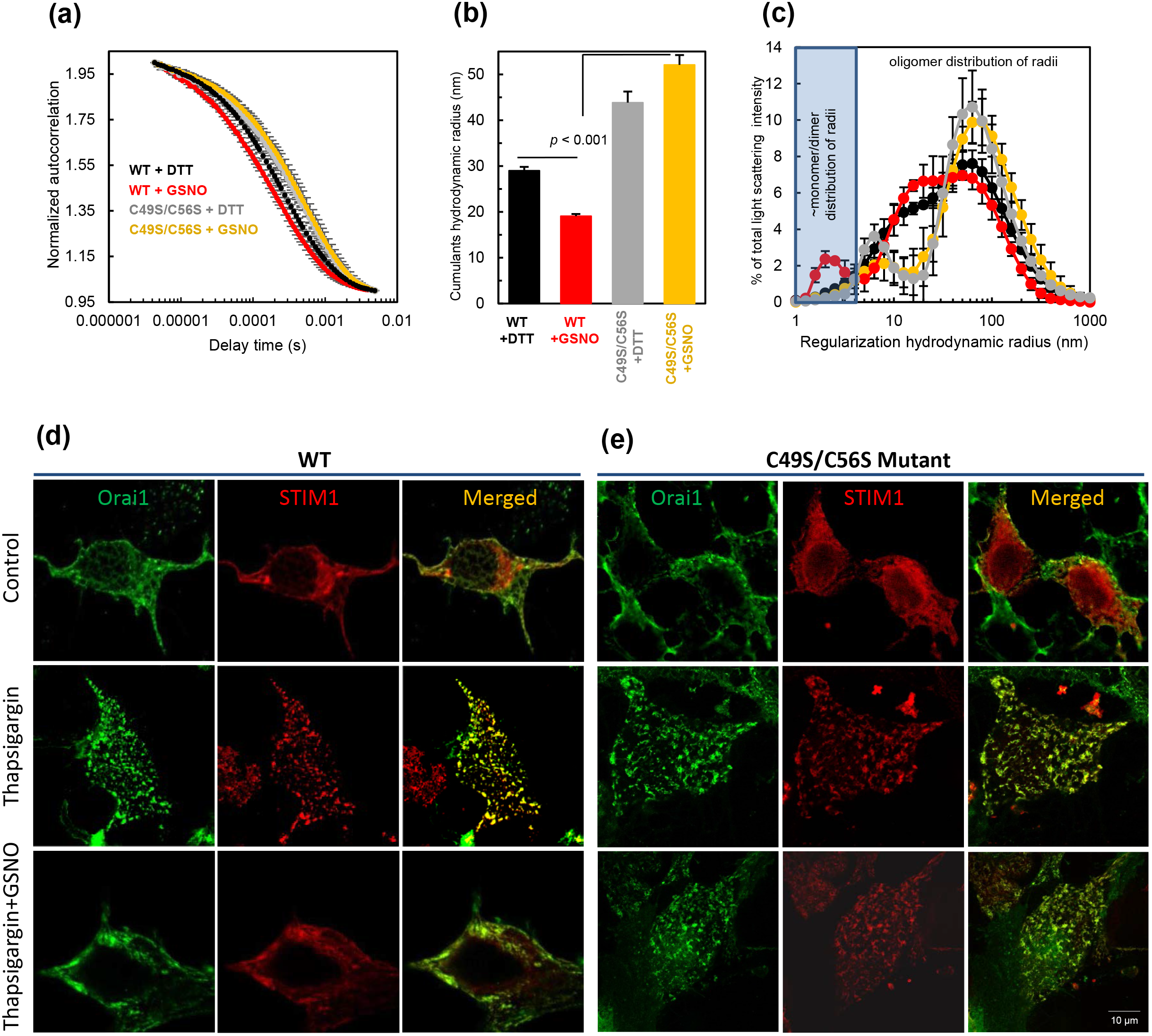
GSNO inhibits STIM1 oligomerization and puncta formation in a Cys-specific manner. **(a-c)** Oligomerization of STIM1 24-213 was examined using dynamic light scattering (DLS). **(a)** Normalized autocorrelation functions of WT and Cys49Ser/C56Ser STIM1 24-213 in the presence of 1 mM DTT or 1 mM GSNO. **(b)** Cumulants fit-derived hydrodynamic radii based on the autocorrelation functions. **(c)** The distribution of hydrodynamic radii for WT + DTT (black) or GSNO (red), Cys49Ser/Cys56Ser mutant + DTT (grey) or GSNO (yellow). Data are means ± SEM from 3 separate protein samples prepared for each condition. **(d-e)** Confocal imaging shows Orai1 (green), STIM1 (red) and merged images of HEK293 cells. Thapsigargin-induced puncta formation of STIM1 was inhibited by GSNO in WT **(d)** but not in Cys49Ser and Cys56Ser mutant STIM1 transfected cells (**e**)

Since the smallest observable radii correspond to the majority of the DLS signal due to the fact that light scattering intensity is proportional to size to the sixth power, this *in vitro* effect is marked [28]. Our data suggest that in the absence of Ca^2+^ and in the presence of the NO donor, oligomerization of the STIM1 luminal domain is suppressed via a mechanism that is dependent on the Cys49 and Cys56 residues.

### NO inhibits mCh-STIM1 puncta formation and co-localization with YFP-Orai1

STIM1 puncta formation and co-localization with Orai1 at SR/ER-plasma membrane junctions is a hallmark of SOCE initiation [29,30]. To assess the effect of NO on STIM1 puncta formation, HEK293 cells stably expressing YFP-Orai1 and transiently expressing WT or Cys49Ser/Cys56Ser mCh-STIM1 were bathed in extracellular Ca^2+^-free medium which contained thapsigargin (2 μM) for 10 min. Confocal imaging revealed that both WT and Cys49Ser/Cys56Ser mCh-STIM1 formed puncta co-localized with YFP-Orai1, indicative of STIM1 activation (**Fig. 4d and e**). Notably, treatment with GSNO (250 μM) prevented puncta formation in WT mCh-STIM1 expressing cells but not in Cys49Ser/Cys56Ser mCh-STIM1 expressing cells (**Fig. 4d and e**). Image quantification using Pearson coefficients shows a significant mCh-STIM1 and YFP-Orai1 colocalization in both WT and double mutant transfected cells after thapsigargin treatment (*P*<0.01, **Fig. S6**). In contrast, incubation with GSNO for 10 min decreased thapsigargin-induced mCh-STIM1 and YFP-Orai1 colocalization in WT (*P*<0.05) but not in double mutant transfected cells (**Fig. S6**). These data are in full agreement with *in vitro* observations showing suppressed oligomerization of S-nitrosylated STIM1 luminal region in a Cys49 and Cys56-dependent manner.

## DISCUSSION

S-Glutathionylation of Cys56 increases the activity of STIM1 and SOCE due to a weakening of STIM1 Ca^2+^ binding affinity [18]. In the present study, the inhibitory effect of S-nitrosylation is opposite to the stimulatory effect of STIM1 S-glutathionylation in non-excitable cells. Further, a propensity for STIM1 Cys49 and Cys56 intramolecular disulfide formation has been previously reported, and blocking this oxidative modification by mutation of the Cys residues to Ala suppresses SOCE in mouse embryonic fibroblasts [20]. This observation is in contrast to data from DT40 cells which showed constitutive SOCE when expressing a STIM1 double Cys49Ala and Cys56Ala mutant [18]. Thus, the phenotype may depend on the cell type used and experimental conditions. Here, we observed a decreased propensity for STIM1 intramolecular disulfide formation in mouse hearts deficient in eNOS and nNOS compared to WT, suggesting a crosstalk between S-nitrosylation and disulfide modifications in cardiomyocytes.

To ascertain the molecular mechanisms underlying the inhibition of STIM1-mediated SOCE by NO, we used recombinant STIM1 proteins corresponding to residues 24-57 and 23-213. Our NMR data revealed that NO modifies Cys49 and Cys56 residues causing structural perturbations associated with decreased mobility of the STIM1 24-57 region. We subsequently showed that the decreased dynamics by NO treatment of the full luminal STIM1 domain encompassing residues 23-213 not only increased the stability of the domain, but also suppressed the oligomerization propensity of the domain in the absence of Ca^2+^. Further, the NO donors did not affect these biophysical properties of the Cys49Ser/Cys56Ser double mutant STIM1 23-213 protein, suggesting that the cysteine residues are critical to STIM1 stabilization. In line with these *in vitro* data we demonstrated that NO inhibits puncta formation of full length mCh-STIM1 in HEK293 cells via a mechanism that is dependent on the Cys49 and Cys56 residues. Collectively, our data suggest a signaling model whereby NO stabilizes the STIM1 protein, inhibits STIM oligomerization and attenuates *I_CRAC_* via S-nitrosylation of Cys49 and Cys56 residues through a structural mechanism that involves decreased mobility of the short nonconserved N-terminal region of the protein. Future high resolution structural data in both the full luminal domain and full STIM1 protein contexts will provide a more complete mechanistic picture of STIM1 regulation by S-NO.

A previous study revealed a consensus I/L-X-C-X_2_-D/E motif which is necessary and sufficient for S-nitrosylation of targets from an inducible NOS (ìNOS)-S100A8-S100A9 complex [31]. However, this motif is not present in STIM1 proteins suggesting an S-nitrosylation mechanism distinct from iNOS and S100 involvement. Nevertheless, we demonstrate that STIM1 S-nitrosylation occurs in WT hearts and is decreased in nNOS^−/−^ and eNOS^−/−^ hearts. While the precise role of eNOS in *I_CRAC_* regulation remains to be determined, we show that nNOS-mediated STIM1 S-nitrosylation inhibits *I_CRAC_* and SOCE in neonatal cardiomyocytes. STIM1 is expressed at significant levels in the neonatal heart and its expression is reduced with a lower SOCE activity in the adult heart but can be induced by cardiac hypertrophy [32]. We now reveal that NO-mediated inhibition of STIM1 activity through S-nitrosylation may also contribute to the lower SOCE in the normal heart. Thus, a functional NO signaling may limit SOCE and cardiac hypertrophy via STIM1 S-nitrosylation. Given that CRAC channels are activated in a graded manner with a steep drop in activity at stoichiometric STIM1:Orai1 binding ratios below 2, not all STIM1 molecules within each channel complex are required to undergo S-nitrosylation to appreciably inhibit CRAC channel activity [33,34]. Under pathological conditions such as heart failure, NO bioavailability is reduced because of higher ROS levels [35,36]. Under these conditions, it is possible that STIM1 S-nitrosylation is decreased while S-glutathionylation is increased, leading to much higher *I_CRAC_* and SOCE, which may contribute to cardiac hypertrophy.

The heart also expresses STIM1L, a splicing variant of STIM1 with the same N-terminal Cys residues and STIM2, which has three Cys residues (Cys15, Cys53 and Cys60) at the luminal region [37,38]. Interestingly, STIM1L activates SOCE immediately upon Ca^2+^ store depletion and allows for repetitive cytosolic Ca^2+^ releases [37]. STIM2 has been shown to mediate increases in resting cytosolic free Ca^2+^ levels in vascular smooth muscle cells (VSMC) and contribute to VSMC proliferation [39]. Additionally, a rare de novo interstitial duplication at 4p15.2, where several candidate genes including STIM2 are located, may potentially cause severe congenital heart defects, limb anomalies, hypogonadism, distinctive facial features, pre- and postnatal developmental delay, and mild cognitive impairments [40]. In the present study, basal intracellular Ca^2+^ levels in neonatal cardiomyocytes are not significantly altered by nNOS. However, we have previously shown that peak Ca^2+^ transient during electric stimulation is increased in nNOS^−/−^ adult mouse cardiomyocytes compared to WT cells [16]. Whether STIM1L and STIM2 can be S-nitrosylated, and how this modification regulates SOCE and intracellular Ca^2+^ levels mediated by these two STIM isoforms in cardiomyocytes remain to be investigated.

In summary, the present study shows that NO can regulate STIM1 activation and SOCE through S-nitrosylation in mammalian cells. Remarkably, our work reveals that the conserved Cys residues on the N-terminal region endow STIM1 with a dynamic functional sensitivity to different thiol modifications depending on the properties of the immediate cellular milieu. Considering the universality of NO and Ca^2+^ signaling in health and disease, we believe the mechanistic and physiologic insights our work provides could open new avenues to the development of STIM1 modulators and greatly expand the understanding of pathophysiological changes signaled via *I_CRAC_* and SOCE.

## METHODS

### Animals

WT and nNOS^−/−^ mice of C57BL/6 background were purchased from The Jackson Laboratory (Bar Harbor, Maine). The investigation conformed to the *Guide for the Care and Use of Laboratory Animals* published by the US National Institute of Health and all experimental protocols were approved by Animal Use Subcommittee at the University of Western Ontario.

### Primary culture of neonatal cardiomyocytes

Neonatal cardiomyocytes were isolated and cultured as described previously [41–43]. Forty-eight hours after seeding, cells were used for Ca^2+^ imaging and patch clamp.

### Ca^2+^ imaging and SOCE measurements

Ca^2+^ imaging was performed using excitation using a deltascan monochrometer system (Photon Technology International, London, ON, Canada) with an automated fluorescence microscope and CCD camera [44]. Neonatal cardiomyocytes were plated on 0.01% polylysine plus 0.1% gelatin-coated Assistant-Präzision coverslips (Deckgläser, Germany). Cardiomyocytes were loaded with 1 μM Fura-2AM (Invitrogen) in extracellular solution (in mM, 140 NaCl, 5 KCl, 1.8 CaCl_2_, 1 MgCl_2_, 10 glucose, and 10 Hepes) for 30 min at room temperature. Next, cells were washed with extracellular solution and kept in this solution until use. Fluorescence intensity at 510 nm was measured with alternating 345 and 380 nm excitation. A standard protocol referred as “Ca^2+^ re-addition” was used to trigger SOCE [45], in which cells were initially exposed to Ca^2+^-free medium containing thapsigargin (2 μM) to deplete SR Ca^2+^ stores. Then, Ca^2+^ (1.8 mM) was restored to the medium, and Ca^2+^ influx was recorded. Data were analyzed using ImageMaster software and shown as 340/380 nm ratios at indicated time points or as time-ratio traces translated from the images.

### Analysis of STIM1 S-nitrosylation in tissues

STIM1 S-nitrosylation was assessed using an S-nitrosylation Biotin Switch Protein Detection Kit (Cayman Chemical, Ann Arbor, MI). Briefly, neonatal hearts from WT and nNOS^−/−^ mice were homogenized in RIPA buffer and lysates were centrifuged at 10,000g to remove unbroken tissues. After protein concentrations were determined by Lowry assay, 500 μg total protein was successively blocked with N-ethylmaleimide, ascorbate reduced and labeled with biotin according to the manufacturer protocol. The free labeling reagents were removed by acetone precipitation, and the pellets were resuspended in 10% HENS buffer/90% neutralization buffer [46]. Biotinylated proteins were pulled out using streptavidin agarose beads and subsequently separated on an 8% SDS-PAGE denaturing gel. S-Nitrosylated STIM1 was indirectly detected as a biotinylated protein after transfer to PVDF membrane by western blotting using a rabbit anti mouse STIM1 antibody (C-terminal; Sigma Aldrich. St. Louis, MO) diluted 1:1500. Aliquots of tissue lysates containing 75 μg of total protein were assessed by western blotting using the same anti-STIM1 antibody as loading controls. To assess STIM1 mobility shift, heart homogenates (50 μg total protein) were run on 5% SDS-PAGE gel under reducing or non-reducing conditions in the presence or absence of 1% β-mercaptoethanol, respectively [20].

### Recombinant protein expression and purification of STIM1 23-213 and 24-57

STIM1 23214 was expressed using a pET-28a vector transformed into BL21 ΔE3 *Escherichia coli* as previously described [6,24]. Briefly, proteins were pulled out of guanidinium solubilized cell lysate using Ni-nitrilotriacetic acid affinity resin (Qiagen). Refolding was done in 20 mM Tris, 300 mM NaCl, 5 mM CaCl_2_, 1 mM DTT, pH 8 by dialysis. The 6×His tags were removed by overnight thrombin digestion (∼1 U mg^−1^ of protein), and the proteins were further purified by Superdex S200 10/300 GL size exclusion chromatography (GE Healthcare). Greater than 95% purity was confirmed by SDS-PAGE and Coomassie blue staining. The Cys mutations were introduced using the QuickChange PCR method (Agilent) and confirmed by sequencing.

Human STIM1 residues 24-57 (NCBI accession NP_003147) was subcloned into the pGEX-4T1 (GE Healthcare) vector using BamHI and EcoRI restriction sites. A Tyr residue was introduced by site-directed mutagenesis immediately N-terminal to residue 24 to simplify protein detection and concentration determinations. The glutathione-S-transferase (GST)-fused STIM1 24-57 protein was expressed in BL21 ΔE3 *E. coli* cells and purified essentially as described in the manufacturer’s GST Sepharose protocol (Genescript). The STIM1 24-57 peptide was liberated from the GST bound to the resin using thrombin (∼ 5 U mg^−1^ of protein). After digestion, the protein was further purified by size exclusion chromatography using the Superdex S200 10/300 GL column.

### Fluorescein labeling of protein S-NO groups

Fifty μL of S-nitrosylated STIM1 23-213 (∼2 mg/mL) protein aliquots were precipitated with two volumes of ice cold acetone followed by centrifugation. The samples were resuspended in 20 mM HEPES, 1 mM EDTA, 2 mM CaCl_2_, pH 7.25 with 200 μM fluorescein-5-maleimide and either 10 mM EDTA or 20 mM ascorbate. After incubation in the dark at 30 °C for 15 min, unreacted fluorescein-5-maleimide was removed by two rounds of acetone precipitation and resuspension in 20 mM HEPES, 2 mM CaCl_2_, pH 7.25. The efficiency of acetone protein precipitation with and without ascorbate was ∼80 %. The precipitated samples were finally resuspended in 600 μL of 20 mM HEPES, 1 mM EDTA, 2 mM CaCl_2_, pH 7.25 and fluorescein fluorescence emission spectra between 500-600 nm were acquired using an excitation wavelength of 494 nm in a Carey Eclipse (Varian/Agilent, Inc.) spectrofluorimeter, temperature equilibrated at 20 °C. The final protein concentration was ∼0.08 mg/mL.

### Solution NMR spectroscopy

Solution nuclear magnetic resonance (NMR) experiments were performed on a 600 MHz Inova NMR spectrometer (Varian/Aglilent, Inc.) equipped with a cryogenic, triple resonance probe. Sequential backbone assignments were made using ^1^H-^15^N HSQC and HNCACB experiments [47]. The two- and three-dimensional spectra were processed, and resonance assignments were made using NMRPipe and XEASY, respectively [48,49].

### Thermal stability by far-UV-CD

Data were acquired on a Jasco J-815 CD Spectrometer (Jasco Inc.) in 1 nm increments (20 nm min^−1^) by using a 0.1 cm pathlength cuvette, an 8-s averaging time and 1 nm bandwidth. Thermal melts were acquired by monitoring the change in the 225 nm CD signal as a function of temperature in 0.1 cm cuvettes, an 8-s averaging time, 1 nm bandwidth and a 1 °C min^−1^ scan rate. The apparent midpoints of temperature denaturation (T_m_) were extracted from the thermal melts using Boltzmann sigmoidal fits in GraphPad.

### Dynamic light scattering

Protein samples were exchanged into condition-specific buffers using centrifugal concentrators (Millipore, Inc.). Proteins samples were centrifuged at 6,000×*g* for 5 min prior to measurement and the upper most portion of the supernatant was immediately taken for the DLS measurements. Protein concentration was 0.36, 0.30 and 0.26 mg mL^−1^ for the WT + DTT samples; 0.38, 0.31 and 0.29 mg mL^−1^ for the WT + GSNO samples; 0.29, 0.22 and 0.11 mg mL^−1^ for the mutant + DTT samples; 0.29, 0.23 and 0.23 mg mL^−1^ for the mutant + GSNO samples. Light scattering measurements were made on a Dynapro DLS module (Protein Solutions, Inc.). The incident light was 825 nm and the scattering angle was 90°. For each sample, 70 consecutive autocorrelation functions, each with a 10 s accumulation time were acquired and the autocorrelation functions from the 10 lowest scattering acquisitions were averaged for hydrodynamic size calculations. The method of Frisken [50] was used for the Cumulants estimation of hydrodynamic radius by fitting 90% of the total decay in the autocorrelation functions. Regularization deconvolution was performed using the accompanying DynaLS software (v1.5) between 4.3 μs – 5.4 ms for all averaged curves using identical resolution settings for each sample. A solvent refractive index of 1.333 and viscosity of 1.019 g m^−1^ s^−1^ was used in the calculations.

### HEK293 cell culture

HEK293 cells stably expressing YFP-Orai1 (generous gift from Dr. Monica Vig, Washington University in St. Louis) were grown in Dulbecco’s Modification of Eagle’s Medium plus 10% fetal bovine serum. The previously described pCMV6 vector encoding full-length human STIM1 (i.e. residues 1-685) with mCh fused just after the ER signal peptide (i.e. residue 1-23) was used in these studies[5]. The Cys49Ser/Cys56Ser mutations were introduced using the QuikChange method. Cells were transfected using an Amaxa Nucleofector following the manufacturer’s guidelines using 0.5 μg mCh-STIM1 or Cys49Ser/Cys56Ser mCh-STIM1. Whole-cell currents were recorded 40-48 h after transfection, and 18-24 h after plating the cells on poly-l-lysine-coated coverslips.

### Electrophysiological Measurement of *I_CRAC_*

Whole-cell voltage-clamp recordings in cultured cardiomyocytes and HEK293 cells were performed at 22-24 °C. The recording electrode was filled with intracellular solution containing (in mM) 145 cesium methane sulfonate, 8 NaCl, 3.5 MgCl_2_, 10 Hepes, and 10 BAPTA (for passive store-depletion and preventing CRAC channel desensitization), pH 7.2. Cells were bathed with extracellular solution consisting of (in mM), 140 NaCl, 5 KCl, 1 MgCl_2_, 10 glucose, and 10 Hepes, pH 7.4, without with Ca^2+^ to prime the CRAC channel proteins, followed by 1.8 CaCl_2_ to induce SOCE. For recordings in cardiomyocytes 0.5 μM tetrodotoxin (a Na^+^ channel blocker) and 10 μM verapamil (a voltage-gated Ca^2+^ channel blocker) were included in the extracellular solution. After incubating cells with extracellular Ca^2+^, transmembrane conductance of test cells (holding voltage = 0 mV) was revealed by a voltage-ramp from −100 to +20 mV in the absence and presence of the CRAC channel inhibitor BTP2 (20 μM). The current-voltage relationship of CRAC channel mediated current (*I_CRAC_*) was displayed by subtracting the current recorded in the presence of BTP2 from the total current recorded in absence of BTP2.

### Confocal Imaging analysis

In order to assess the impact of GSNO on oligomerization of STIM1, confocal imaging analysis was employed. Briefly, HEK293 cells stably expressing YFP-Orai1 were transfected with full length WT mCh-STIM1 or Cys49Ser/Cys56Ser mCh-STIM1. Fluorescence cell images were captured using Zeiss LSM 510 laser confocal microscope under control conditions and after Ca^2+^ depletion and thapsigargin treatment with and without GSNO for 10 minutes. Colocalization of mCh-STIM1 and YFP-Orai1 was analyzed using Image J (version 1.51, NIH) to calculate Pearson correlation coefficients.

### Statistical Analysis

All data are expressed as mean ± SEM. One-way analysis of variance (ANOVA) followed by Tukey’s post hoc test was used to compare differences among groups. Differences were considered statistically significant at *P*<0.05.

## Author Contributions

L.G. performed the experiments, analyzed the data and wrote the first draft of the manuscript. J.Z. performed NMR and thermal stability experiments, and analyzed the data. X.L. and S.M.S. help with experimental design and data analysis. W.Y.L., P.B.S. and Q.F. deigned experiments, and revised and edited the manuscript.

## ACKNOWLEDGEMENTS

This study was supported by operating grants from Canadian Institutes of Health Research (MOP-142383 to Q.F., P.B.S. and W.Y.L.), the Natural Sciences and Engineering Research Council of Canada (05239 to P.B.S) and National Natural Science Foundation of China (NSFC-81570294 to L.G.). L.G. was a visiting scholar from Nantong University, Nantong, China.

## REFERENCES

[1] L. Majewski, J. Kuznicki, SOCE in neurons: Signaling or just refilling?, Biochim Biophys Acta. 1853 (2015) 1940–52.

[2] J.A. Stiber, P.B. Rosenberg, The role of store-operated calcium influx in skeletal muscle signaling, Cell Calcium. 49 (2011) 341–9.

[3] H.E. Collins, X. Zhu-Mauldin, R.B. Marchase, J.C. Chatham, STIM1/Orai1-mediated SOCE: current perspectives and potential roles in cardiac function and pathology, Am J Physiol Heart Circ Physiol. 305 (2013) H446–58.

[4] P.B. Stathopulos, M. Ikura, Partial unfolding and oligomerization of stromal interaction molecules as an initiation mechanism of store operated calcium entry, Biochem Cell Biol. 88 (2010) 175–83.

[5] P.B. Stathopulos, L. Zheng, G.Y. Li, M.J. Plevin, M. Ikura, Structural and mechanistic insights into STIM1-mediated initiation of store-operated calcium entry, Cell. 135 (2008) 110–22.

[6] P.B. Stathopulos, G.Y. Li, M.J. Plevin, J.B. Ames, M. Ikura, Stored Ca^2+^ depletion-induced oligomerization of stromal interaction molecule 1 (STIM1) via the EF-SAM region: An initiation mechanism for capacitive Ca^2+^ entry, J Biol Chem. 281 (2006) 35855–62.

[7] R.M. Luik, B. Wang, M. Prakriya, M.M. Wu, R.S. Lewis, Oligomerization of STIM1 couples ER calcium depletion to CRAC channel activation, Nature. 454 (2008) 538–42.

[8] J. Liou, M. Fivaz, T. Inoue, T. Meyer, Live-cell imaging reveals sequential oligomerization and local plasma membrane targeting of stromal interaction molecule 1 after Ca^2+^ store depletion, Proc Natl Acad Sci USA. 104 (2007) 9301–6.

[9] M.M. Wu, J. Buchanan, R.M. Luik, R.S. Lewis, Ca^2+^ store depletion causes STIM1 to accumulate in ER regions closely associated with the plasma membrane, J Cell Biol. 174 (2006) 803–13.

[10] J. Soboloff, B.S. Rothberg, M. Madesh, D.L. Gill, STIM proteins: dynamic calcium signal transducers, Nat Rev Mol Cell Biol. 13 (2012) 549–65.

[11] X. Zhu-Mauldin, S.A. Marsh, L. Zou, R.B. Marchase, J.C. Chatham, Modification of STIM1 by O-linked N-acetylglucosamine (O-GlcNAc) attenuates store-operated calcium entry in neonatal cardiomyocytes, J Biol Chem. 287 (2012) 39094–106.

[12] E. Pozo-Guisado, D.G. Campbell, M. Deak, A. Alvarez-Barrientos, N.A. Morrice, I.S. Alvarez, et al., Phosphorylation of STIM1 at ERK1/2 target sites modulates store-operated calcium entry, J Cell Sci. 123 (2010) 3084–93.

[13] Y.J. Choi, Y. Zhao, M. Bhattacharya, P.B. Stathopulos, Structural perturbations induced by Asn131 and Asn171 glycosylation converge within the EFSAM core and enhance stromal interaction molecule-1 mediated store operated calcium entry, Biochiw Biophys Acta. 1864 (2017) 1054–1063.

[14] S.M. Haldar, J.S. Stamler, S-nitrosylation: integrator of cardiovascular performance and oxygen delivery, J Clin Invest. 123 (2013) 101–10.

[15] D.R. Gonzalez, F. Beigi, A.V. Treuer, J.M. Hare, Deficient ryanodine receptor S-nitrosylation increases sarcoplasmic reticulum calcium leak and arrhythmogenesis in cardiomyocytes, Proc Natl Acad Sci USA. 104 (2007) 20612–7.

[16] D.E. Burger, X. Lu, M. Lei, F.L. Xiang, L. Hammoud, M. Jiang, et al., Neuronal nitric oxide synthase protects against myocardial infarction-induced ventricular arrhythmia and mortality in mice, Circulation. 120 (2009) 1345–1354.

[17] P.B. Stathopulos, M. Ikura, Structure and function of endoplasmic reticulum STIM calcium sensors, Curr Top Membr. 71 (2013) 59–93.

[18] B.J. Hawkins, K.M. Irrinki, K. Mallilankaraman, Y.C. Lien, Y. Wang, C.D. Bhanumathy, et al., S-glutathionylation activates STIM1 and alters mitochondrial homeostasis, J Cell Biol. 190 (2010) 391–405.

[19] D.R. Arnelle, J.S. Stamler, NO^+^, NO, and NO^−^ donation by S-nitrosothiols: implications for regulation of physiological functions by S-nitrosylation and acceleration of disulfide formation, Arch Biochem Biophys. 318 (1995) 279–85.

[20] D. Prins, J. Groenendyk, N. Touret, M. Michalak, Modulation of STIM1 and capacitative Ca^2+^ entry by the endoplasmic reticulum luminal oxidoreductase ERp57, EMBO Rep. 12 (2011) 1182–8.

[21] Y. Miao, C. Miner, L. Zhang, P.I. Hanson, A. Dani, M. Vig, An essential and NSF independent role for alpha-SNAP in store-operated calcium entry, eLIFE. 2 (2013) e00802.

[22] N.E. Hafsa, D. Arndt, D.S. Wishart, CSI 3.0: a web server for identifying secondary and super-secondary structure in proteins using NMR chemical shifts, Nucleic Acids Res. 43 (2015) W370–7.

[23] M.V. Berjanskii, D.S. Wishart, A simple method to predict protein flexibility using secondary chemical shifts, J Am Chem Soc. 127 (2005) 14970–1.

[24] P.B. Stathopulos, L. Zheng, M. Ikura, Stromal interaction molecule (STIM) 1 and STIM2 calcium sensing regions exhibit distinct unfolding and oligomerization kinetics, J Biol Chem. 284 (2009) 728–32.

[25] E.D. Covington, M.M. Wu, R.S. Lewis, Essential role for the CRAC activation domain in store-dependent oligomerization of STIM1, Mol Biol Cell. 21 (2010) 1897–907.

[26] M. Muik, M. Fahrner, R. Schindl, P. Stathopulos, I. Frischauf, I. Derler, et al., STIM1 couples to ORAI1 via an intramolecular transition into an extended conformation, EMBO J. 30 (2011) 1678–89.

[27] Y. Zhou, P. Srinivasan, S. Razavi, S. Seymour, P. Meraner, A. Gudlur, et al., Initial activation of STIM1, the regulator of store-operated calcium entry, Nat Struct Mol Biol. 20 (2013) 973–81.

[28] Y. Li, V. Lubchenko, P.G. Vekilov, The use of dynamic light scattering and brownian microscopy to characterize protein aggregation, Rev Sci Instrum. 82 (2011) 053106.

[29] P.B. Stathopulos, M. Ikura, Structurally delineating stromal interaction molecules as the endoplasmic reticulum calcium sensors and regulators of calcium release-activated calcium entry, Immunol Rev. 231 (2009) 113–31.

[30] S. Perni, J.L. Dynes, A.V. Yeromin, M.D. Cahalan, C. Franzini-Armstrong, Nanoscale patterning of STIM1 and Orai1 during store-operated Ca^2+^ entry, Proc Natl Acad Sci U S A. 112 (2015) E5533–42.

[31] J. Jia, A. Arif, F. Terenzi, B. Willard, E.F. Plow, S.L. Hazen, et al., Target-selective protein S-nitrosylation by sequence motif recognition, Cell. 159 (2014) 623–34.

[32] X. Luo, B. Hojayev, N. Jiang, Z.V. Wang, S. Tandan, A. Rakalin, et al., STIM1-dependent store-operated Ca^2+^ entry is required for pathological cardiac hypertrophy, J Mol Cell Cardiol. 52 (2012) 136–47.

[33] Z. Li, L. Liu, Y. Deng, W. Ji, W. Du, P. Xu, et al., Graded activation of CRAC channel by binding of different numbers of STIM1 to Orai1 subunits, Cell Res. 21 (2011) 305–15.

[34] P.J. Hoover, R.S. Lewis, Stoichiometric requirements for trapping and gating of Ca^2+^ release-activated Ca^2+^ (CRAC) channels by stromal interaction molecule 1 (STIM1), Proc Natl Acad Sci USA. 108 (2011) 13299–304.

[35] G.K. Couto, L.R. Britto, J.G. Mill, L.V. Rossoni, Enhanced nitric oxide bioavailability in coronary arteries prevents the onset of heart failure in rats with myocardial infarction, J Mol Cell Cardiol. 86 (2015) 110–20.

[36] U. Forstermann, W.C. Sessa, Nitric oxide synthases: regulation and function, Eur Heart J. 33 (2012) 829–37, 837a-837d.

[37] B. Darbellay, S. Arnaudeau, C.R. Bader, S. Konig, L. Bernheim, STIM1L is a new actin-binding splice variant involved in fast repetitive Ca^2+^ release, J Cell Biol. 194 (2011) 335–46.

[38] P.B. Stathopulos, M. Ikura, Structural aspects of calcium-release activated calcium channel function, Channels. 7 (2013) 344–53.

[39] S. Song, S.G. Carr, K.M. McDermott, M. Rodriguez, A. Babicheva, A. Balistrieri, et al., STIM2 (Stromal Interaction Molecule 2)-Mediated Increase in Resting Cytosolic Free Ca(2+) Concentration Stimulates PASMC Proliferation in Pulmonary Arterial Hypertension, Hypertension. 71 (2018) 518–529.

[40] L. Liang, Y. Xie, Y. Shen, Q. Yin, H. Yuan, A Rare de novo Interstitial Duplication at 4p15.2 in a Boy with Severe Congenital Heart Defects, Limb Anomalies, Hypogonadism, and Global Developmental Delay, Cytogenet Genome Res. 150 (2016) 112–117.

[41] W. Song, X. Lu, Q. Feng, Tumor necrosis factor-alpha induces apoptosis via inducible nitric oxide synthase in neonatal mouse cardiomyocytes, Cardiovasc. Res. 45 (2000) 595–602.

[42] T. Zhang, X. Lu, P. Arnold, Y. Liu, R. Baliga, H. Huang, et al., Mitogen-activated protein kinase phosphatase-1 inhibits myocardial TNF-alpha expression and improves cardiac function during endotoxemia, Cardiovasc Res. 93 (2012) 471–9.

[43] Y. Wu, C. Qin, X. Lu, J. Marchiori, Q. Feng, North American ginseng inhibits myocardial NOX2-ERK1/2 signaling and tumor necrosis factor-alpha expression in endotoxemia, Pharmacol Res. 111 (2016) 217–225.

[44] X. Luo, D.M. Shin, X. Wang, S.F. Konieczny, S. Muallem, Aberrant localization of intracellular organelles, Ca^2+^ signaling, and exocytosis in Mist1 null mice, J Biol Chem. 280 (2005) 12668–75.

[45] G.S. Bird, W.I. DeHaven, J.T. Smyth, J.W. Putney, Jr., Methods for studying store-operated calcium entry, Methods. 46 (2008) 204–12.

[46] M.T. Forrester, M.W. Foster, M. Benhar, J.S. Stamler, Detection of protein S-nitrosylation with the biotin-switch technique, Free Radic Biol Med. 46 (2009) 119–26.

[47] S. Grzesiek, A. Bax, Amino acid type determination in the sequential assignment procedure of uniformly 13C/15N-enriched proteins, J Biomol NMR. 3 (1993) 185–204.

[48] C. Bartels, T.H. Xia, M. Billeter, P. Guntert, K. Wuthrich, The program XEASY for computer-supported NMR spectral analysis of biological macromolecules, J Biomol NMR. 6 (1995) 1–10.

[49] F. Delaglio, S. Grzesiek, G.W. Vuister, G. Zhu, J. Pfeifer, A. Bax, NMRPipe: a multidimensional spectral processing system based on UNIX pipes, J Biomol NMR. 6 (1995) 277–93.

[50] B.J. Frisken, Revisiting the method of cumulants for the analysis of dynamic light-scattering data, Appl Opt. 40 (2001) 4087–91.

